# Chitin utilization by marine picocyanobacteria and the evolution of a planktonic lifestyle

**DOI:** 10.1101/2022.06.23.497379

**Authors:** Giovanna Capovilla, Rogier Braakman, Gregory Fournier, Thomas Hackl, Julia Schwartzman, Xinda Lu, Alexis Yelton, Krista Longnecker, Melissa Kido Soule, Elaina Thomas, Gretchen Swarr, Alessandro Mongera, Jack Payette, Jacob Waldbauer, Elizabeth B. Kujawinski, Otto X. Cordero, Sallie W. Chisholm

## Abstract

Marine picocyanobacteria (*Prochlorococcus* and *Synechococcus*), the most abundant photosynthetic cells in the oceans, are generally thought to have a primarily single-celled and free-living lifestyle. However, we find that genes for breaking down chitin - an abundant source of organic carbon that primarily exists as particles - are widespread in this group. We further show that cells with a chitin degradation pathway display chitin degradation activity, attach to chitin particles and show enhanced growth under low light conditions when exposed to chitosan, a partially deacetylated form of chitin. Marine chitin is largely derived from arthropods, whose roots lie in the early Phanerozoic, 520-535 million years ago, close to when marine picocyanobacteria began colonizing the ocean. We postulate that attachment to chitin particles allowed benthic cyanobacteria to emulate their mat-based lifestyle in the water column, initiating their expansion into the open ocean, seeding the rise of modern marine ecosystems. Transitioning to a constitutive planktonic life without chitin associations along a major early branch within the *Prochlorococcus* tree led to cellular and genomic streamlining. Our work highlights how coevolution across trophic levels creates metabolic opportunities and drives biospheric expansions.

---

*Prochlorococcus* and its sister lineage marine *Synechococcus* (together ‘marine picocyanobacteria’) are a monophyletic and highly abundant group of oceanic phytoplankton that perform about ∼25% of oceanic CO_2_-fixation(1–3). In addition to their role as primary producers, it is increasingly clear that marine picocyanobacteria also use organic carbon as a supplemental carbon and energy source (i.e. mixotrophy)(4–6), especially in the light-limited deep euphotic zone(7, 8). While studying genes involved in mixotrophy in this group(9), we noticed that some strains have chitinase genes, indicating the potential for using chitin, one of the most abundant sources of organic carbon in the ocean(10, 11). However, chitin in the marine environment typically exists as particles, primarily derived from arthropod exoskeletons(10, 11), which are broken down by microbial consortia(12). Marine picocyanobacteria are typically considered to live an exclusively single-celled planktonic lifestyle(13, 14), but are also sometimes found within particulate organic matter aggregates(15) that sink out of the euphotic zone(16). Together this suggests that chitin utilization might be a functional trait in marine picocyanobacteria.

To explore this possibility we began by examining the distribution of genes involved in chitin utilization in *Prochlorococcus* and *Synechococcus* genomes (Fig. 1). This comparison is complicated by the significant overlap in gene content between pathways for chitin utilization and peptidoglycan recycling (Fig. 1). Peptidoglycan, which is chemically similar to chitin, forms a single cell-size polymeric structure (the sacculus) that is a component of the bacterial cell wall, and recycling of peptidoglycan fragments inevitably generated during cell growth and division is widespread in bacteria(17). We therefore searched 702 partial and complete genomes for genes required for both chitin utilization and peptidoglycan recycling. We found that in *Synechococcus* and deeply branching “low-light-adapted IV” (LLIV) *Prochlorococcus* most of these genes are nearly universal, i.e. they occur at a frequency similar to the average genome completeness of ∼75% (Methods) (Fig. 1). In contrast, these genes are almost universally absent in “high-light-adapted” (HL) *Prochlorococcus* (Fig. 1). All LL clades of *Prochlorococcus* possess peptidoglycan recycling genes, but chitin utilization genes are nearly exclusive to the deeply branching LLIV clade (Fig. 1). These observations suggest that chitin utilization and peptidoglycan recycling were both present in the last common ancestor of marine picocyanobacteria, but were lost during the diversification of extant *Prochlorococcus* groups.

**Figure 1.**
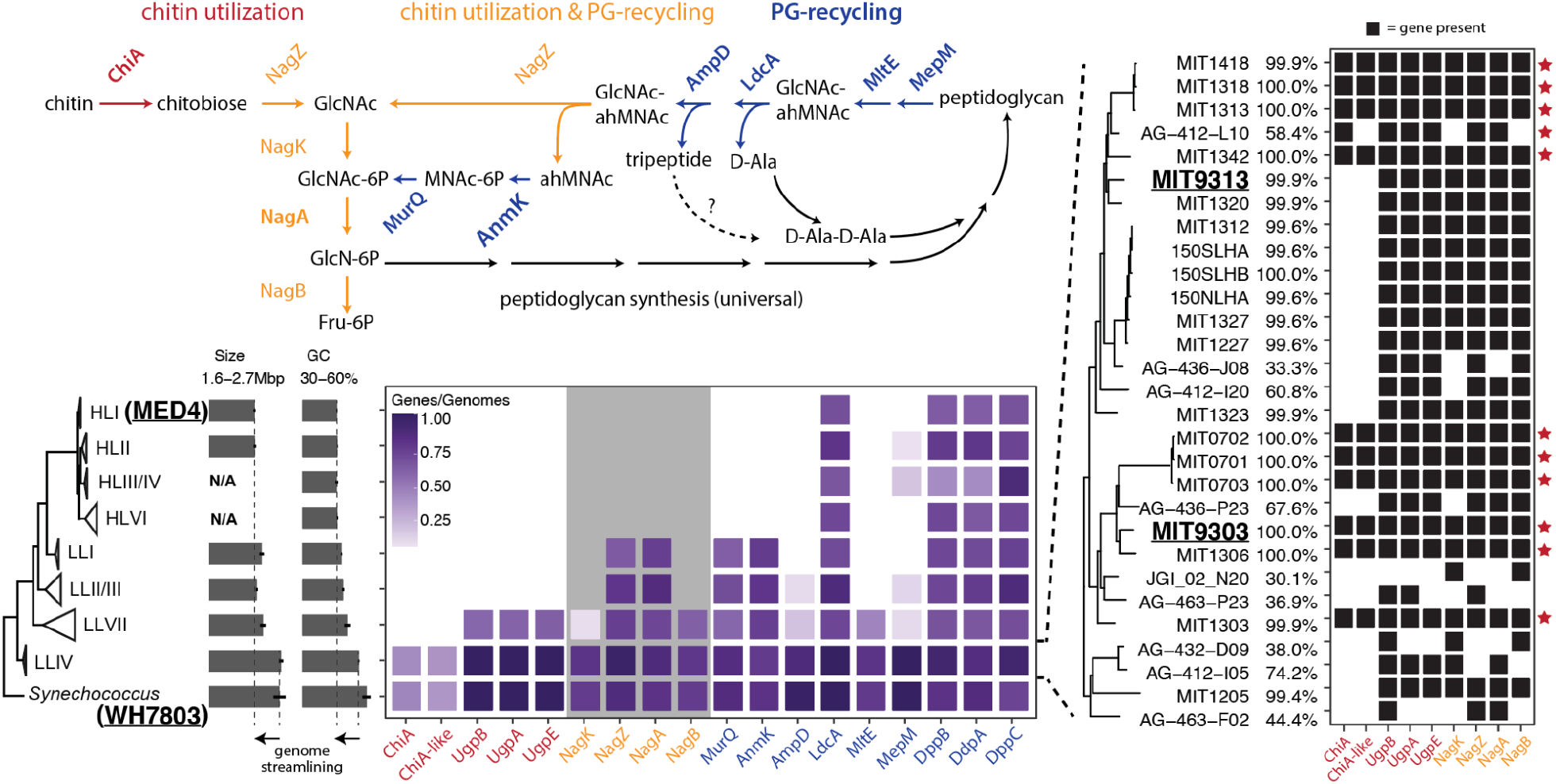
Distribution of chitin utilization genes in marine picocyanobacteria. <u>Upper panel</u>: Pathway for chitin degradation (red) and its reactions that overlap (orange) with peptidoglycan metabolic recycling (blue). ChiA is annotated as a putative chitinase enzyme, while ChiA-like is a homolog of the substrate-binding domain of ChiA and annotated as a putative chitin-binding domain protein. <u>Lower panel</u>: Average frequency of occurrence of chitin utilization and peptidoglycan recycling genes estimated from both partial and complete genome sequences available in *Synechococcus* and the major clades of *Prochlorococcus* shown in the tree on the left. Average completeness of genomes in our sample is ∼75% (see Methods). The clade-membership of the strains used in the experiments are highlighted in bold and underlined. The grey background area in the gene frequency frame highlights genes shared between the chitin utilization and peptidoglycan pathways (orange). The panel on the right breaks down members of the LLIV clade of *Prochlorococcus*, revealing putative primary chitin degraders of chitin that possess chitinase (red star) and putative secondary degraders that lack chitinase. Abbreviations: GlcNAc - N-acetyl-glucosamine, GlcNAc-6P - N-acetyl-glucosamine 6-phosphate, GlcN-6P - glucosamine 6-phosphate, F6P - fructose 6-phosphate, ahMNAc - anhydro-N-acetyl-beta-muramate, MNAc-6P - N-acetyl-muramate 6 phosphate.

In both *Prochlorococcus* and *Synechococcus*, genomes containing chitin-utilization genes sub-differentiate into two groups, which either possess or lack chitinase, the enzyme that hydrolyzes chitin(Fig. 1). It is known from other systems that chitin utilization is a complex ecological process(18, 19) with hydrolysis of chitin polymers occurring in the extracellular milieu, making chitin fragments available to all cells in the community. Consequently, chitin breakdown involves niche partitioning into groups that specialize in performing the initial hydrolysis steps and others that specialize in using small chitin oligosaccharides(20). Our results suggest a similar niche partitioning into ‘primary’ and ‘secondary’ degraders may occur among chitin-using members of the marine picocyanobacteria.

To determine whether the putative chitin users (Fig. 1) actually metabolize chitin, we first used enzyme assays to examine whether the cells display chitinase activity. Both *Synechococcus* and *Prochlorococcus* strains that are putative primary degraders of chitin (i.e. possessing the full complement of chitin utilization genes) displayed chitinase activity in cell-free spent media, and the activity disappeared upon boiling the media (Fig. 2, S1) – consistent with the production of extracellular chitinase enzymes that are denatured upon heating. In agreement with the genomic predictions, a *Prochlorococcus* strain (MIT9313, Fig. 1) that is a putative secondary chitin degrader did not display extracellular chitinase activity in spent media (Fig. 2, S1).

**Figure 2.**
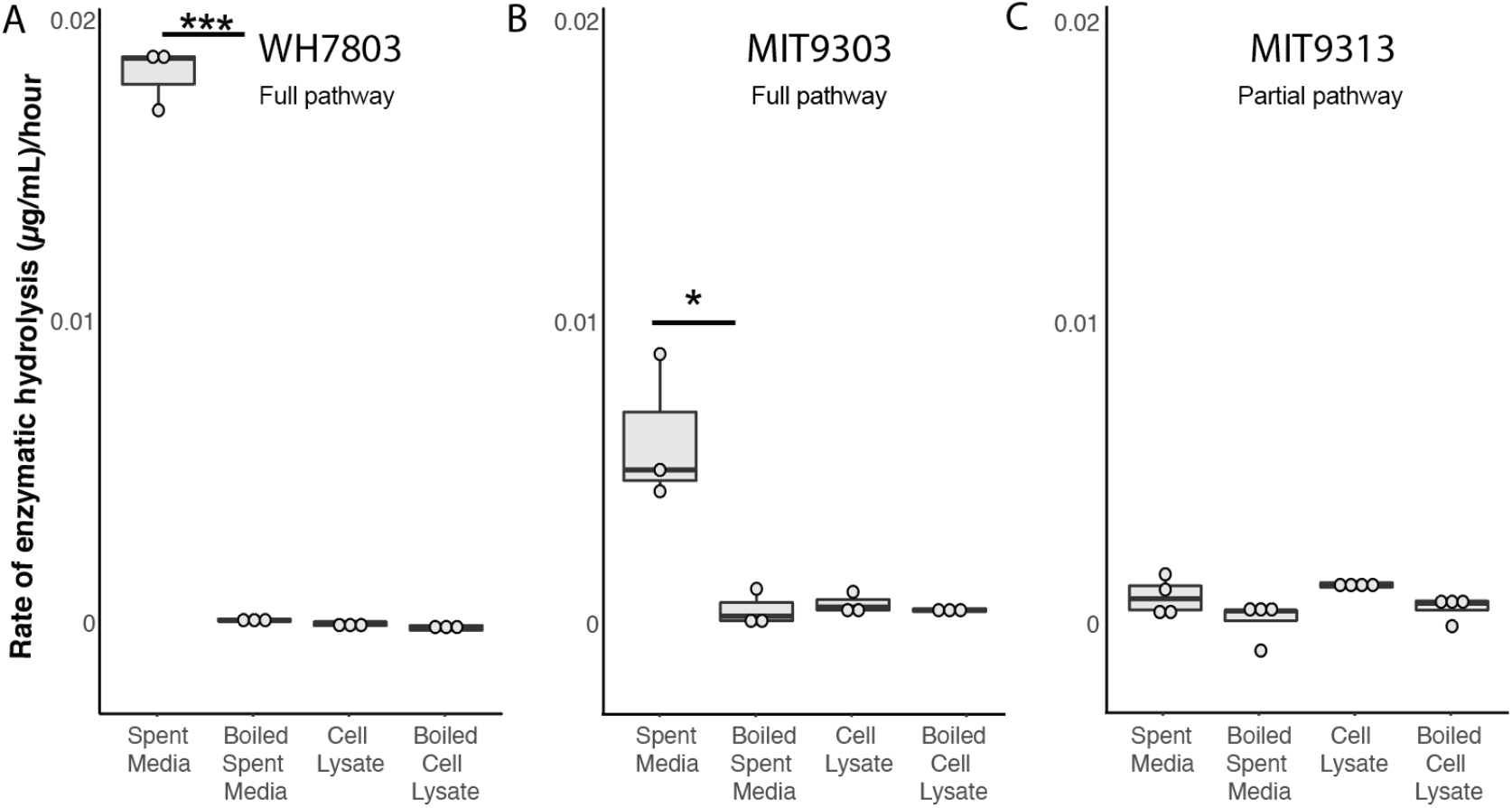
Chitinase activity in *Prochlorococcus* and *Synechococcus*. Endochitinase activity in *Synechococcus* WH7803, *Prochlorococcus* MIT9303, and *Prochlorococcus* MIT9313 measured in spent media and cell lysates. Strains with the complete chitin degradation pathway display such activity in the spent media. Activity is lost after boiling. Exochitinase activity is shown in Figure S1.

The next question we addressed was whether or not *Synechococcus* and *Prochlorococcus* cells with the chitin utilization pathway colonize chitin surfaces, by adding hydrogel chitin beads to cultures. The *Prochlorococcus* strains used were from the two sub-groups we identified as putative primary (MIT9303) and secondary (MIT9313) degraders (Fig. 1), as well as cells lacking the complete pathway for chitin degradation (MED4). The *Synechococcus* strain used was WH7803, a primary chitin degrader with a complete set of chitin utilization genes. We found that both *Synechococcus* and *Prochlorococcus* cells with chitin utilization genes attach to chitin particles, while *Prochlorococcus* lacking chitin degradation genes do not (Fig. 3). Primary degraders attach both to the surface of particles and accumulate in “pockets” within the beads (Fig. 3A,C,D,F), while secondary degraders only accumulate in pockets (Fig. 3A,B,F), potentially reflecting the inferred niche partitioning between them. Finally, we found that *Prochlorococcus* attaches only to chitin particles and not to agarose, but that *Synechococcus* attaches to both (Fig. 3A, S2), suggesting that *Synechococcus* has broader surface attachment abilities than *Prochlorococcus*.

**Figure 3.**
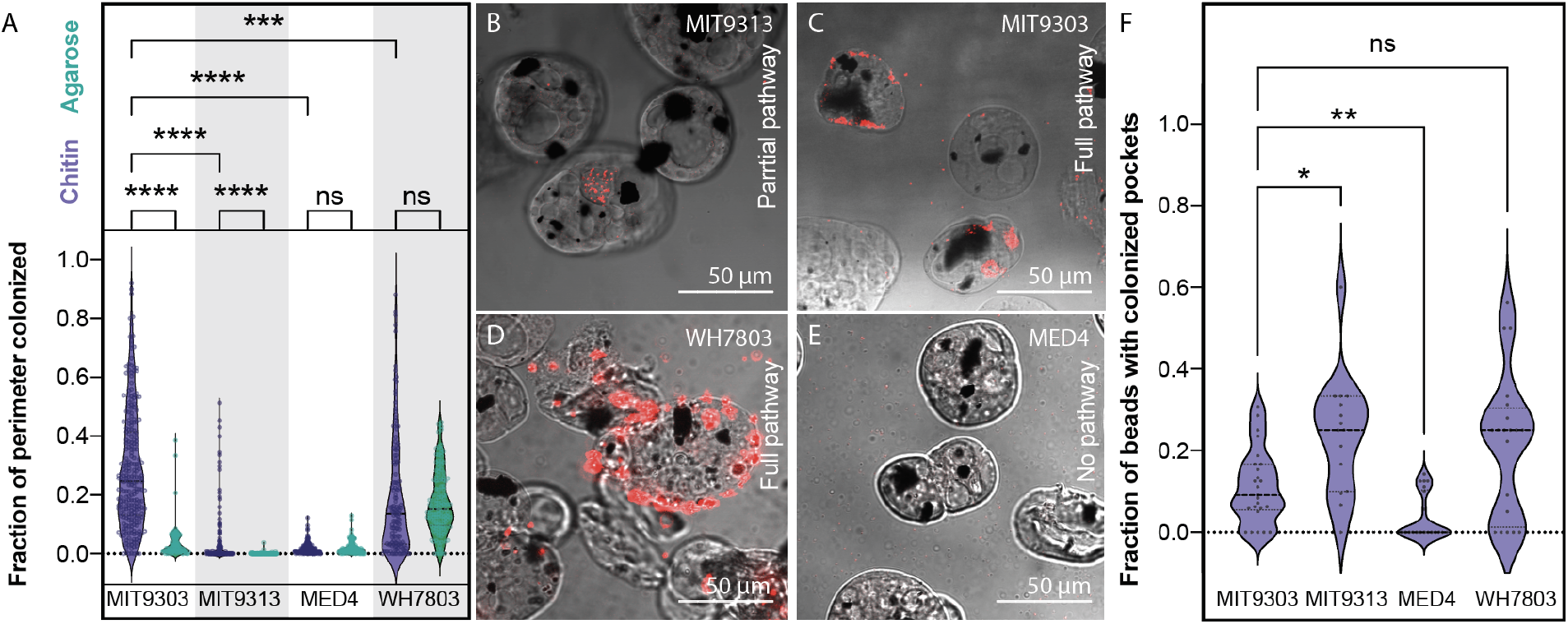
Colonization of artificial chitin particles by *Prochlorococcus* and *Synechococcus*. A) Fraction of particle perimeter colonized by a strain of *Synechococcus* (WH7803) and 3 strains of *Prochlorococcus* (MED4, MIT 9313, and MIT 9303) exposed to agarose (green) and chitin (purple) particles. *Synechococcus* WH7803 and *Prochlorococcus* MIT9303 are putative primary chitin degraders, *Prochlorococcus* MIT9313 is a putative secondary chitin degrader (i.e. it lacks chitinase), and *Prochlorococcus* MED4 lacks all chitin degradation genes. B-E). Confocal sections of chitin particles quantified in A. Chitin particles are shown in bright field mode; the dark spots are magnetic beads embedded in the particles. *Synechococcus* and *Prochlorococcus* are detected by their red autofluorescence. Cells were sometimes observed contained in pockets within the particles (as shown in B and C, for example). Quantification of this phenomenon is shown in F.

As a material resource, chitin potentially provides energy, carbon, and/or nitrogen. *Prochlorococcus* niche partitioning provides relevant clues on what it derives from chitin, as chitin utilization is only retained within the LLIV clade (Fig. 1), which dominates at the bottom of the euphotic zone(21). At these depths, light goes to extinction and *Prochlorococcus* is thought to derive a significant fraction of its carbon through mixotrophy rather than photosynthesis(7, 8). This suggests chitin could provide *Prochlorococcus* a supplemental carbon and energy source that is beneficial under low light conditions. To test this hypothesis, we grew *Prochlorococcus* cells representing both the primary and secondary degrader genotypes (Fig. 1) at several light levels, with and without the addition of chitosan, a form of chitin that is solubilized through partial deacetylation. Differences due to chitosan additions emerged only at the lowest light level (Fig. 4), and for primary degraders was expressed as a higher growth rate, albeit not significantly so, while for secondary degraders was expressed as a higher cell density in stationary phase, potentially reflecting their niche partitioning. A growth boost due to chitin utilization only under light-limitation is consistent with results in other cyanobacteria showing that wild-type cells able to recycle peptidoglycan fragments generated during the normal course of growth have higher growth rates than mutants without this ability, but only at low light levels(22). We also examined whether *Prochlorococcus* can use chitosan as a source of nitrogen by adding it to the growth media of nitrogen-starved cells, but relief from nitrogen stress was not apparent (Fig. S3, S4). Together, this suggests that in its role as a resource, chitin primarily acts as a supplemental carbon source to light-limited *Prochlorococcus* cells deep in the water column.

**Figure 4.**
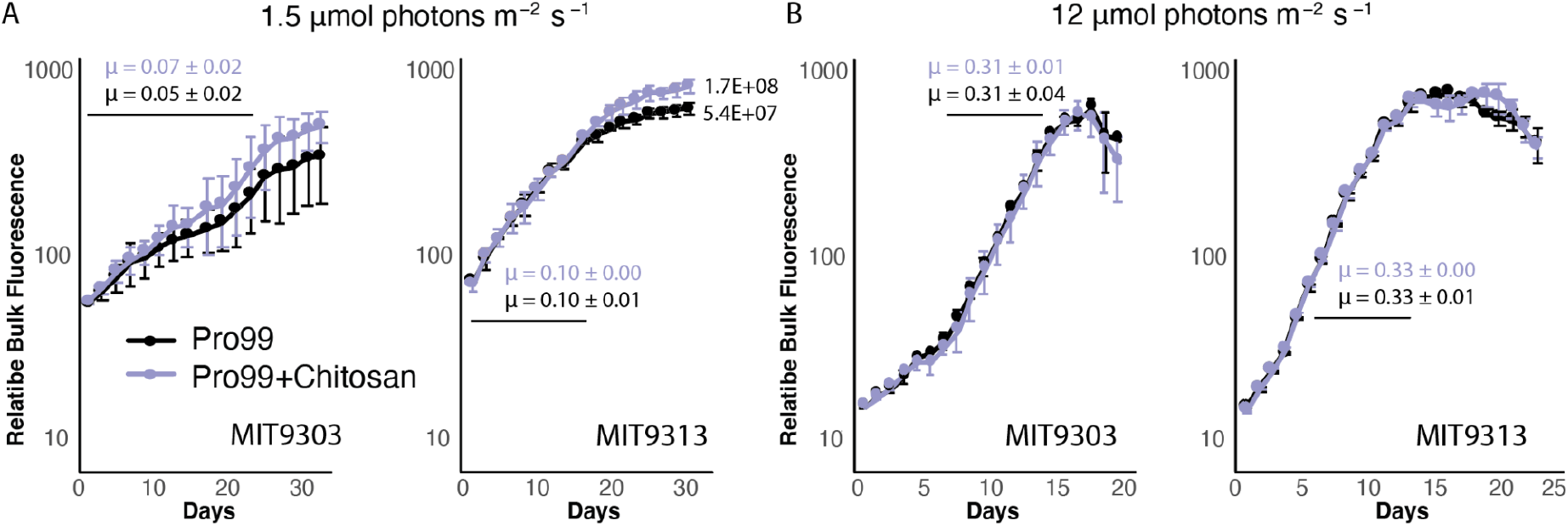
Effect of chitosan addition on growth of two *Prochlorococcus* strains as a function of light intensity. MIT9303 is a primary chitinase degrader and MIT9313 is a secondary degrader, missing the chitinase genes. Cultures were grown in Pro99 media with (purple) and without (black) added chitosan, at two different light intensities: 1.5 μmol photons m^-1^ s^-1^ (A) and 12 μmol photons m^-1^ s^-1^ (B), the former being growth-limiting for these strains as reflected in the difference in growth rates, μ, at the different light levels. Growth was monitored by bulk chlorophyll fluorescence, but upon emergence of a significant difference in fluorescence in MIT9313 cultures cell counts were measured using flow cytometry. Chitosan-amended cultures had higher cell counts in stationary phase (1.7E+8 cells mL^-1^) than unamended cultures (5.4E+7 cells mL^-1^), reflecting an increase in cell yield. Error bars show standard deviation between the 3 biological replicates. The average growth rate and associated standard deviation (μ, in units day^-1^) was calculated in exponential phase (marked with a black line) and is shown for each curve.

To investigate how cells respond to chitin addition at the molecular level, we used qPCR to measure the expression of genes in the chitin degradation pathway in *Prochlorococcus* cultures grown with and without chitosan. All genes were expressed under both conditions (Table S3), including the chitinase and chitobiose transporter genes that are specific to chitin degradation (Fig. 5A). This suggests cells constitutively express this pathway, perhaps so they are poised to use chitin when it appears in the environment. Furthermore, while changes in expression due to chitosan exposure were not statistically significant for most genes, we observed a clear trend across the full suite of genes. That is, shortly after chitosan addition, expression of chitinase genes was higher relative to unamended cultures, while expression of downstream genes was lower (Fig. 5A). At later time points, the expression of chitinase was lower and downstream genes higher in amended relative to unamended cultures (Fig. 5A). These patterns suggest that *Prochlorococcus* cells increase the activity of chitin degradation enzymes in the order in which pathway intermediates become available, which is broadly similar to how chitin degradation pathways operate in other chitinolytic bacteria(23).

**Figure 5.**
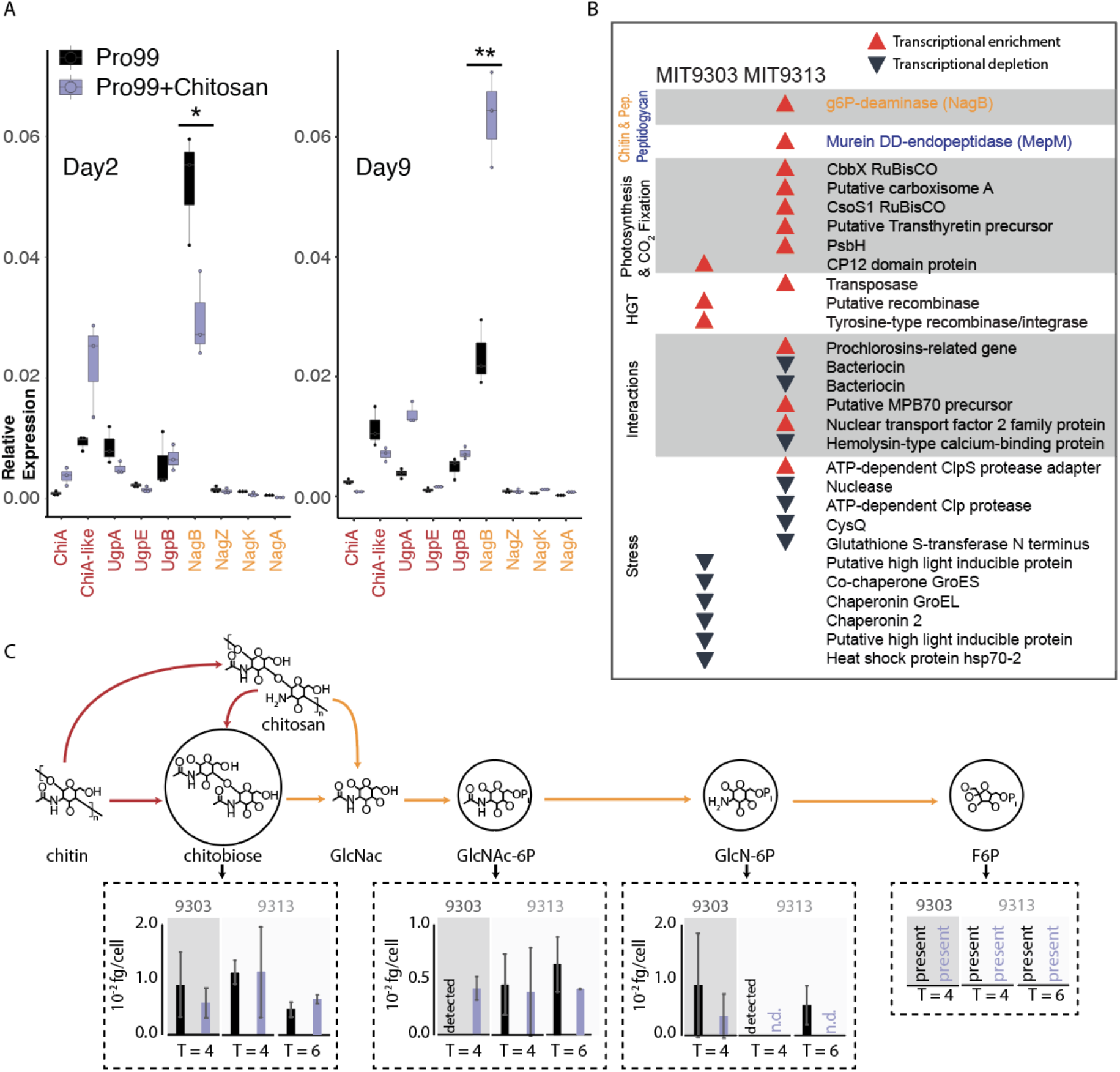
Gene expression and metabolomic analysis in *Prochlorococcus* in response to the addition of chitosan. A) Expression (measured by qPCR) of all the chitin-related genes in MIT9303 (primary degrader) in relation to the housekeeping gene, *rnpB*, gene in natural seawater-based Pro99 medium in presence and absence of chitosan at two time points over the growth curve. Cells were in early exponential growth on Day 2 and mid-exponential on Day 9. B) Qualitative representation of the relative expression of genes (measured using RNA-Seq) in the two *Prochlorococcus* strains 24 hours after addition to chitosan (see also Fig. S5). Red upward pointing arrows represent statistically significant transcript enrichment in presence of chitosan while black downward pointing arrows represent statistically significant transcript depletion. Genes are grouped by functional categories, which are shown in shaded/unshaded sections. C) Concentrations of intermediates of chitin degradation on days 4 and 6 after chitosan additions (purple) compared to unamended controls (black). Error bars show the standard deviations of 3 biological replicates for MIT9303 and MIT9313 on day 4, and of 2 replicates for MIT9313 on day 6. Molecular abbreviations and color scheme is the same as in Fig. 1. Metabolites are labeled as ‘detected’ in samples where they were observed in only a single replicate, and as ‘n.d.’ (not detected) in samples where they were not observed in any replicates. Fructose-6-P was present in most samples, but its levels could not be quantified due to interference by the organic carbon matrix.

To augment the qPCR results and gain a systems-level understanding of how the availability of chitin shapes *Prochlorococcus*, we performed RNA-Seq and metabolomic analyses of cultures exposed to chitosan. While expression of many genes changed in response to the addition of chitosan, relatively few of those changes were significant, and they disappeared after 48 hours (Fig. 5B, S5). As with our analysis of the expression of individual genes by qPCR, this suggests a mild and transient response to chitosan additions. We further observed transcripts of all genes of the chitin utilization pathway and accumulation of all its metabolic intermediates under all conditions (Fig. 5C, Table S3,S4), consistent with the idea that cells constitutively express this pathway to be ready when chitin appears in the environment, as also suggested by qPCR data (Fig. 5A).

While the chitin utilization pathway is always active, *Prochlorococcus* metabolism also responds to chitosan additions: in one strain, MIT9313 (secondary degrader), the expression of two genes involved in peptidoglycan recycling (NagB, MepM), one of which is also involved in chitin utilization (NagB), significantly increased (Fig. 5B), consistent with the noted molecular, physiological and ecological links between pathways (Fig. 1). Similarly, concentrations of chitin degradation intermediates also changed upon addition of chitosan, although most changes were not significant (Fig. 3C). Additional changes occurred in downstream metabolic processes. Transcripts of several genes involved in photosynthesis and carbon fixation were enriched in the presence of chitosan in MIT9313, a secondary degrader of chitin, which could have indirectly contributed to the difference in cell number observed in low light levels (Fig. 4A). Transcripts for CP12, which inhibits carbon fixation(24), were enriched in the presence of chitosan in MIT9303, a primary degrader of chitin (Fig. 1,3B). Furthermore, the concentration of several intermediates of core carbohydrate metabolism changed in the presence of chitosan (Fig S6), although again few changes were significant. Together these observations indicate that chitosan exposure modifies the processing of carbon through core metabolic pathways.

Changes in a number of other functional categories in cells exposed to chitosan can be grouped together in terms of their inferred ecological function. Transcripts of various genes involved in horizontal gene transfer (e.g., transposases and recombinases), and microbial interactions (e.g., antibiotics, lanthipeptides, secreted proteins) were enriched, while transcripts of various genes involved in mitigating stress (e.g., chaperonins, high light-inducible proteins, proteases, and genes related to glutathione) were depleted (Fig. 3B, S5). Since life in biofilms increases microbial interactions, promotes horizontal gene transfer(25), helps buffer against external stresses(26), and increases likelihood of the light-limited conditions under which peptidoglycan recycling becomes beneficial(22), we interpret these collective changes as cells preparing to attach to chitin particles and switch to a surface-attached lifestyle.

To gain insight into the Earth-historical context of the acquisition of the chitin utilization trait in *Prochlorococcus* and *Synechococcus*, we examined when this acquisition happened relative to the divergence of marine picocyanobacteria from other cyanobacteria. Picocyanobacterial sequences for most genes involved in both the chitin degradation and peptidoglycan recycling pathways (i.e. orange genes in Fig. 1) are nested within larger branches of cyanobacterial genes (Fig. 6, S7), indicating ancestral vertical inheritance of peptidoglycan recycling. In contrast, picocyanobacterial sequences for both chitinase (ChiA and ChiA-like) and N-acetylglucosamine kinase (NagK) are nested within non-cyanobacterial diversity, indicating horizontal gene transfer to picocyanobacteria after their divergence from other cyanobacteria (Fig. 6, S7). NagK is present in deeply branching ‘SynPro’ groups (i.e., including *Cyanobium* and related *Synechococcus* that bridge the fresh-salt water divide), but chitinase genes are exclusive to ‘marine SynPro’ (Fig. 6, S7). NagK fortifies peptidoglycan recycling by completing a second branch involved in metabolizing GlcNAc but is obligately required for chitin utilization (Fig. 1,7A). These observations suggest that peptidoglycan recycling pre-dated the radiation of marine picocyanobacteria and provided a preadaptation to chitin utilization. Acquisition of chitinase along the stem leading from total group SynPro to crown-group marine SynPro then enabled chitin utilization by this group (Fig. 7A).

**Figure 6.**
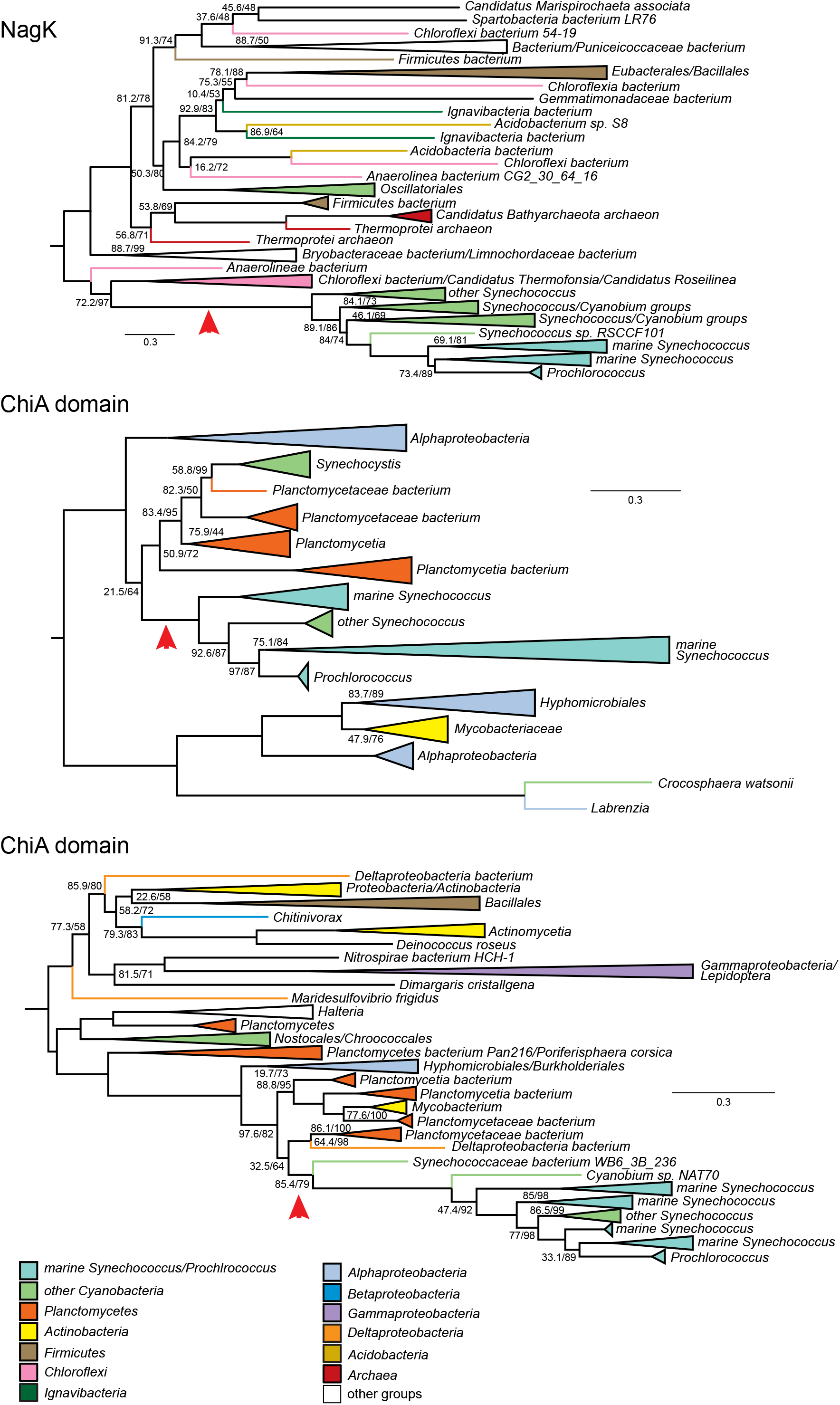
Phylogenies of ChiA and NagK genes in marine picocyanobacteria. Red arrows indicate inferred HGTs into SynPro ancestor lineages. Branch lengths are indicated by included scale bars (average substitutions/site). High-level taxonomic identities are indicated by color coding, with specific represented groups labeled for terminal and collapsed groups. In each case, trees are rooted at the branch closest to the midpoint that preserves the monophyly of Synechococcales groups including SynPro. Branch supports (approximate likelihood ratio test/bootstrap value) are included for bipartitions with less than 90% support for either metric. Phylogenies of NagZ, NagA, NagB, UgpA, and UgpB are found in Figure S7.

**Figure 7.**
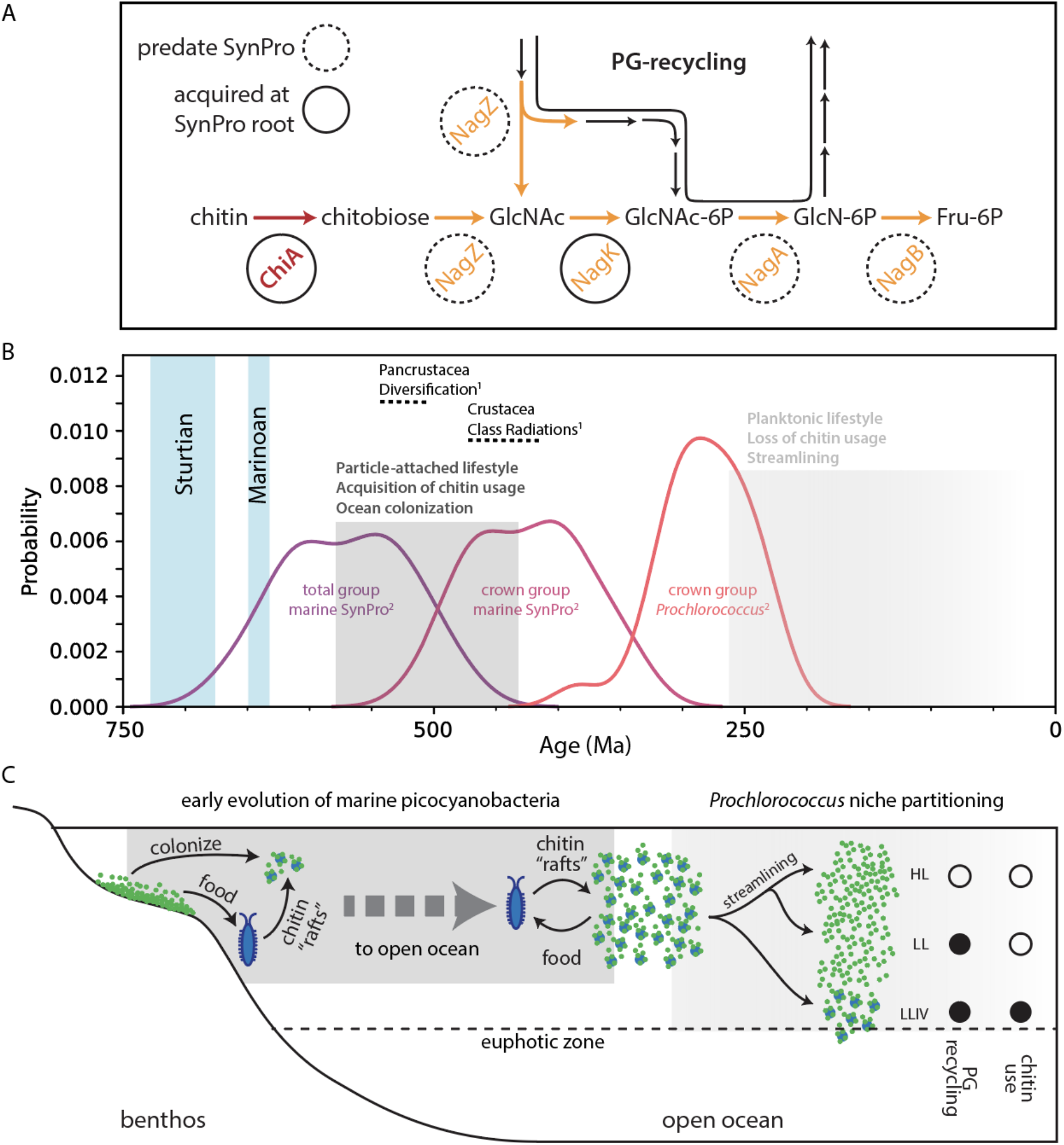
Hypothesized timing and ecological context of the rise of marine picocyanobacteria. A) Pathways for chitin utilization and peptidoglycan metabolic recycling. Genes exclusive to chitin utilization are shown in red, and genes shared between the two pathways are shown in orange. Genes highlighted with dashed circles pre-date evolution of marine picocyanobacteria, while genes highlighted with solid circles were acquired along the stem leading to their crown group. B) Summary of divergence time estimates for the evolution of 1) arthropods(27) and 2) marine picocyanobacteria(28), which we postulate are linked via chitin utilization metabolism. Vertical blue bars represent the snowball Earth intervals preceding the rise of marine picocyanobacteria. C) The “chitin-raft hypothesis” for the rise of marine picocyanobacteria. The rise of marine arthropods led to an accumulation of chitin in the environment, providing a substrate that benthic cyanobacteria could colonize while preserving their surface-attached lifestyle. Existing peptidoglycan recycling pathways provided a preadaptation for acquiring chitin degradation pathways (Fig. 7A). As *Prochlorococcus* diversified, loss of chitin associations and adaptation to a constitutive planktonic lifestyle occurred during cellular and genomic streamlining, and a shift from physical-chemical interactions within particle-attached communities to purely chemical interactions in the water column. A facultative ability to colonize chitin particles and use chitin as a supplemental carbon source under light limitation was preserved in the LLIV clade of *Prochlorococcus* that is adapted to the low light conditions at the bottom of the euphotic zone.

To better understand the context of the transfer of chitinase genes into ancestors of marine SynPro we examined their phylogenetic relationships to similar sequences found within other bacterial groups. Picocyanobacterial chitinases contain two major chitin-binding domains that show homology to different chitinase sequence variants found within other bacterial genomes. Gene sequence alignments suggest that the marine SynPro variant is likely the product of a fusion of two genes that were both acquired from Planctomycetes (SI Text 1). However, the sampling of Planctomycete diversity is too sparse to determine if the gene fusion event occurred before or after the transfers into ancestral marine SynPro (see SI Text 1 and SI Text 1 files for detailed discussion). Planctomycete genomes and metagenomes containing chitinases are found in both fresh and marine environments, preventing us from determining if the gene transfers to picocyanobacteria occurred in freshwater environments prior to colonization of the ocean, or within the marine planktonic environment. Recipients of chitinase genes from Planctomycetes include both marine and freshwater bacteria (SI Text 1), suggesting the lack of a strong congruence between ecological and phylogenetic signals.

Palaeobiological evidence provides further insight into the early evolution of chitin utilization in marine picocyanobacteria. First, molecular clocks constrained by horizontally transferred genes that link the timing of cyanobacterial evolution to that of other bacterial phyla indicate that the stem lineage leading to crown group SynPro existed during the interval of ∼570-420 Ma(28) (Fig. 7B). Second, both fossils and fossil-calibrated molecular clocks indicate that marine arthropods – whose exoskeletons are the main source of chitin in the ocean(10, 11) – underwent a major ecological expansion between ∼535-400 Ma(27, 29, 30). Finally, while the body fossil record from open ocean environments is limited because of a high likelihood of degradation before preservation, the second half of this period (∼480-400 Ma), nearer to the calculated timing of crown group SynPro (Fig. 7B), is thought to have seen an increase in pelagic planktonic arthropods(31) and the emergence of more complex trophic structures in the open ocean(31–34). These dates are consistent with the hypothesis that the early evolution of marine picocyanobacteria and arthropods was intertwined, and played a role in establishing modern marine ecosystems.

Given chitin utilization by marine picocyanobacteria and their apparent contemporaneous evolution with arthropods, one might expect existence of direct associations with arthropods in the extant biosphere. However, while *Synechococcus* has been found to be abundant in copepod guts (35), we found no reports of direct associations with the exoskeletons of living copepods or other arthropods(35, 36). It is possible that chitin utilization by picocyanobacteria has become primarily restricted to detrital particulate matter, derived from exoskeletal molts (which constitute the largest flux of arthropod chitin into the environment) and dead bodies. Alternatively, direct arthropod-picocyanobacterial associations may yet be discovered now that there is added motivation to look for them. Indeed, picocyanobacterial symbionts have been discovered in association with an increasingly diverse set of algae in recent years, including dinoflagellates, foraminifera, radiolarians, and tintinnids(15, 37–41).

The emergence of chitin attachment and utilization during the rise of marine picocyanobacteria helps address a conundrum regarding the evolution of lifestyle in this group. In most environments other than lakes and oceans, microbes predominantly live as aggregates attached to surfaces, which provides protection from external stresses and allows efficient recycling of nutrients(26). The earliest marine cyanobacteria likely also predominantly lived in mats in the benthic environment(13, 42–44). In contrast, extant marine picocyanobacteria are generally thought to live a single-celled planktonic life(13, 14), despite existing in a saline, UV-rich, and nutrient-poor environment, which are all factors associated with inducing aggregation of cells and attachment to surfaces(26, 45–47). This raises the question: what was the sequence of innovations that allowed cyanobacteria to make the evolutionary transition from life in benthic mats to life as individual planktonic cells in the open ocean?

To resolve this conundrum, we propose the “chitin raft hypothesis”, in which picocyanobacteria and arthropods colonized the open ocean in tandem (Fig. 7C). Accumulated detrital chitin and other organic material could have provided “rafts” that allowed picocyanobacteria to maintain a surface-attached lifestyle while expanding into the ocean. By promoting collective nutrient recycling and stress mitigation, life in chitin particle-attached communities would have provided refugia from the harsh environment of the open ocean, allowing cells over millions of years of evolution to acquire necessary adaptations for an eventual transition to a single-celled planktonic life. Indeed, evolution of marine picocyanobacteria involved acquisition of genes for synthesis of unique compatible solutes for mitigating osmotic stress in saline environments(48, 49), various genes involved in mitigating light/UV stress or repairing light damage(50, 51), a number of changes to the membrane, genome, and proteome that are thought to lower cellular nutrient requirements(52–54), and a general remodeling of metabolism that is thought to enhance cellular nutrient affinity(55). Each of these innovations acts to mitigate against stresses that in other contexts are linked to triggering aggregation of cells and attachment to surfaces(26, 45– 47).

This sequence of innovations culminated with a period of dramatic genomic and proteomic streamlining(52, 54) (Fig. 1) and enhanced genetic drift(56) along the branch separating the LLIV clade and all other *Prochlorococcus*, indicating the occurrence of a major population bottleneck. This transition also involved a shift in other *Prochlorococcus* lineages toward a smaller and rounder cell(57, 58) and the loss of both chitin utilization genes (Fig. 1) and the ability to attach to chitin particles (eg. MED4 in Fig. 3). Together these observations lead us to conclude that the divergence between LLIV and other *Prochlorococcus* involved a transition from a facultatively particle-attached to a constitutively planktonic lifestyle. Metabolic reconstructions indicate that streamlining was part of a drive toward greater cellular energy flux with as a by-product increased organic carbon exudation, in turn driving co-evolution with co-occurring heterotrophs(55). Added context from our findings here suggests that various adaptations over the course of *Prochlorococcus* evolution collectively mediated an ecological transition from physical-chemical interactions at microscopic scales on particles to primarily chemical interactions at ecosystem scales in the water column.

## Materials and Methods

### Culture conditions for growth curves

*Prochlorococcus* or *Synechococcus* cells were grown under constant light flux at 24°C in natural seawater-based Pro99 medium containing 0.2-μm-filtered Sargasso Sea water, amended with Pro99 nutrients (N, P, and trace metals) prepared as previously described (Moore *et al*. 2007). The low light experiment in *Prochlorococcus* MIT9303 and MIT9313 was performed at a light level of 1.5 μmol quanta m^−2^ s^−1^. The other experiments in *Prochlorococcus* or *Synechococcus* were performed at 12 or 15 μmol quanta m^−2^ s^−1^ as specified in each legend. The final concentrations of NH_4_^+^were 800 μM in Pro99, 250 μM in Pro5, 150 μM in Pro3, and 50 μM in Pro1. In these experiments, half of the samples were amended with high molecular weight chitosan (Millipore Sigma) to a final concentration of 56 μg/ml. All the growth curves were performed in triplicates.

### Identification of genes involved in chitin degradation and peptidoglycan recycling

To identify genes potentially involved in chitin-degradation, we searched for homologs to known chitin degradation genes(12) in *Prochlorococcus* strains MIT9313 and MIT1318 using blast. Similarly, for peptidoglycan recycling, we searched for homologs to known peptidoglycan recycling genes(17). We found homologues for most genes of the peptidoglycan recycling pathway employed by *E. coli(17)* in *Prochlorococcus*, most of which had previous functional annotations consistent with our findings. We next assigned each of these sequences to NCBI COGs (clusters of orthologous groups)(59) based on reciprocal best hits (Table S1_chitin-cogs)

We then expanded these annotations to our entire collection: 623 *Prochlorococcus(60)* and 79 *Synechococcus* genomes(61) (Table S1_pro-syn-genomes). Of note, the majority of these genomes were derived from single-cell sequencing projects and often are not complete. The estimated average completeness for genomes in this set is 75%. We first clustered all proteins using mmseqs(62) (--cov-mode 5 -c 0.5), then annotated the cluster representatives with eggNOG-mapper(63) to obtain NCBI COG labels for each cluster, and, finally, crossmatched the clusters with the COG-labels assigned to our seed sequence set. We manually checked and if necessary refined each matched protein cluster for consistency and correct functional assignment. For the final list of annotated genes see (Table S1 pro-chitin-genes).

### Quantitative PCR analysis

*Prochlorococcus* MIT9303 cells grown at 11 μmol photons m^−2^ s^−1^ were collected from the samples by centrifugation in triplicates at each time point and each condition. RNA samples were extracted with a standard acidic Phenol:Chloroform protocol and measured with Nanodrop (Thermo Scientific). RevertAid First Strand cDNA Synthesis Kit (Thermo Scientific) with random primers was used to obtain cDNA. Quantitative PCR reactions were performed in a CFX96 thermocycler (Bio-Rad) using the primers listed in Table S1. The expression of *rnpB* gene was used to normalize the results.

### Chitinase assay

*Prochlorococcus* MIT9303, MIT9313 and *Synechococcus* WH7803 cultures were grown at 15 μmol quanta m^−2^ s^−1^ in Pro99 media amended with high molecular weight chitosan (Millipore Sigma) to a final concentration of 56 μg/ml in triplicates. A volume of 50 ml of culture in mid-exponential was then centrifuged to separate the cells fraction and the spent media. The pellet was flash frozen, resuspended in sterile MilliQ water to a volume of 2 ml, sonicated and filtered through a Spin-X centrifuge tube filter (Costar). The supernatant was filtered through a 0.45 μm filter and concentrated using 30kDa Amicon® Ultra-15 Centrifugal Filter Units (Millipore) to a volume of 1,5 ml. Half volume of each sample has been boiled at 90°C for 45 minutes to serve as controls. Protease inhibitor (Roche) was added to all samples. Each sample was then divided into 3 aliquots. Each aliquot was tested with one of the 3 substrates contained in the Chitinase kit (Sigma): 4-Methylumbelliferyl N,N′- diacetyl-β-D-chitobioside (substrate suitable for exochitinase activity detection or chitobiosidase activity), 4-Methylumbelliferyl N-acetyl-β-D-glucosaminide (substrate suitable for exochitinase activity detection or β-N-acetylglucosaminidase activity) and 4- Methylumbelliferyl β-D-N,N′,N′′-triacetylchitotriose (substrate suitable for endochitinase activity detection).

The aliquots were then mixed with each of these substrates, incubated in darkness and the fluorescence of the 4-methyl-umbelliferone released by the chitinase activity in the sample was measured every 24 hours on a plate reader set at excitation 360 nm and emission at 450 nm.

### Beads and microscopy

Chitin magnetic beads (NEB Cat#E8036L), or Agarose magnetic beads (Pierce™ Glutathione Magnetic Agarose Beads Thermo Scientific™ Cat#78602) were washed in Pro99 media and size selected using cell strainers. Chitin beads smaller than 40 μm or larger than 100 μm have been discarded during the washes with Pro99. The media enriched with beads has then been inoculated with the *Prochlorococcus* or *Synechococcus* strains of interest, in triplicates in light level of 30 μmol quanta m^−2^ s^−1^ (for MED4 and WH7803) and 11 μmol quanta m^−2^ s^−1^ (for MIT9303 and MIT9313). Growth was monitored using bulk culture fluorescence measured with a 10AU fluorometer (Turner Designs) and samples were collected in late exponential, at a cell density of 10^8. A volume of 1ml of each sample was poured on a 35mm Petri dish with 20 mm Glass Bottom Microwell Dish (Mat Tek). To avoid excessive fluorescent signal from floating cyanobacteria, the samples were washed 5 times with fresh media. Floating culture was removed by placing the mini dish on a magnet (to keep the magnetic beads on the bottom) and then substituted with fresh Pro99 media before placing the sample on the scope. Images were taken with a LSM780 Zeiss, capturing the beads on the bright field and the *Prochlorococcus* by its autofluorescence with laser 561.

### Analysis of the colonized perimeter and cells colonizing pockets

To quantify the fraction of cells colonizing the perimeter of chitin or agarose magnetic beads, we wrote a custom analysis script using MATLAB (version R2020a, Mathworks. Code available at https://github.com/jaschwartzman/prochlorococcus). The script segmented the brightfield image of magnetic beads to create a binary mask for the bead perimeter. To account for the fact that chitin magnetic beads are semi-transparent, the mask was created using the gradient of the image, which defines the shadow around the perimeter of the beads. This area was filled and eroded to define the bead center. The center area was subtracted from the total area to define the perimeter zone of the beads. Any cells found in this zone are assumed to be attached to the bead surface. To define cell area, we segmented the accompanying image of cell autofluorescence. Briefly, cells were segmented by defining a linear threshold based on the magnitude of the gradient of the autofluorescence image. This threshold was defined for each day of experimental acquisition, to account for variation in acquisition parameters. For each bead, the perimeter area, and the subset of the perimeter area colonized by cells were quantified. The fraction of the perimeter area colonized was defined by dividing the total perimeter area by the cell area. An assumption of this quantification is that cells from different taxa occupy approximately the same 2-dimensional area.

The chitin magnetic beads were found to contain pockets, in which we sometimes observed cells. The irregularities and pockets internal to the chitin hydrogel were visible due to the weak autofluorescence of chitin detected in the 561 nm channel and were also identifiable in brightfield images. In the latter scenario, the greater depth of field in the brightfield image creates the possibility that cells are sitting on a surface above a hollow feature. Due to the diverse morphologies of pockets, we were unable to define a consistent set of characteristics to segment these structures using a script. Accordingly, we quantified the number of particles in which we observed cells colonizing internal pockets manually, by counting instances of cells colonizing either a zone with less autofluorescence within a particle and verifying the presence of a pocket in the brightfield image.

### RNA-Seq and Metabolites analysis

*Prochlorococcus* MIT9303 and MIT9313 cells were grown under constant light flux at 11 μmol photons m^−2^ s^−1^ at 24°C in artificial AMP1 medium(64). Cultures were transferred to AMP1 from Pro99 and kept in AMP1 for at least 3 transfers before performing the experiment to avoid any potential contributions to observed pathway dynamics by background levels of uncharacterized organic polymers in natural seawater. Half of the samples were supplemented with chitosan (Millipore Sigma) to a final concentration of 56 μg/ml. The growth curves were monitored using bulk culture fluorescence measured with a 10AU fluorometer (Turner Designs) and the cell count was recorded using Guava® easyCyte™ HT Flow Cytometer. Samples were collected on day 1 and day 3 after inoculation for RNA extraction and day 4 and day 6 for the metabolomics analysis.

For the RNA-Seq *Prochlorococcus* cells were collected from the samples by centrifugation in triplicates at each time point and each condition. RNA samples were extracted with a standard acidic Phenol:Chloroform protocol, rRNA was depleted with NEBNext^®^ rRNA Depletion Kit (Bacteria); libraries were prepared with KAPA HyperPrep Kit (Roche) and the multiplexed samples were run on an Illumina HiSeq with a 75nt NextSeq sequencing program.

### RNA-Seq analysis

The raw Illumina reads were trimmed of adapters using bbduk v38.16(65) (ktrim=r, k=23, mink=11, hdist=1). Low-quality regions were removed from the adapter-trimmed sequences with bbduk v38.16(65) (qtrim=rl, trimq=6). The trimmed RNA-seq reads from *Prochlorococcus* MIT9303 and MIT9313 cultures were aligned to the MIT9303 and MIT9313 reference genomes, respectively (available from https://github.com/thackl/protycheposons/) with the Burrows-Wheeler Aligner v0.7.16a-r1181(66), using the BWA-backtrack algorithm. The number of reads that aligned to each annotated ORF in the “sense” orientation was determined using the HTSeq package v0.11.2(67) (default parameters, “nonunique all”). Counts of reads that aligned to each ORF (excluding rRNAs and tRNAs) were compiled across replicates. MIT9303 and MIT9313 reads were analyzed separately using the DESeq2 R package v1.24.0(68) to determine differentially expressed genes. The standard DESeq2 functions and workflow were implemented to normalize samples by library sequencing depth and estimate gene dispersion. Differential expression tests were performed on comparisons of cultures with chitosan vs. control cultures one and three days after chitosan addition. Significance was determined with the Wald test, using a negative binomial generalized linear model, and p-values were corrected for multiple testing with the Benjamini–Hochberg procedure. Following the guidance of the DESeq2 authors(68), genes were considered to be significantly differentially expressed between a given pair of treatments if the adjusted p-value was <0.1. Gene expression results were visualized with ggplot2(69).

### Metabolite extractions

For the metabolomics analysis, *Prochlorococcus* cells were filtered under gentle vacuum (to prevent cell lysis) in a glass filtration tower onto 0.1 μM Omnipore filters (Millipore). Filters were then folded and placed into a cryovial, and frozen at −80°C until processed. We extracted intracellular metabolites using a method described in (70) and modified as described in (71). Briefly, we extracted each filter with ice-cold extraction solvent (40:40:20 acetonitrile: methanol: water with 0.1 M formic acid), neutralized the combined extracts with 6M ammonium hydroxide, and dried them in a vacufuge until near dryness. Prior to liquid chromatography-mass spectrometry (LC/MS) analysis, we re-constituted all extracts in 245 μL of solvent (30 μL MQ pure water, 70 μL methanol, and 145 μL acetonitrile) with isotopically-labeled injection standards (50 pg/μL each; d_2_ biotin, d_6_ succinic acid, d_4_ cholic acid, d_7_ indole-3-acetic acid, and ^13^C_1_-phenylalanine). We generated a pooled sample for quality control (QC) by combining 25 μL aliquots from each sample.

### LC/MS analysis of metabolites

The organic matter extracts were analyzed using targeted LC/MS. A mix of metabolite standards (see Table S4) was made in 80:20 acetonitrile:Milli-Q water at 10 μg/mL. Matrix-matched calibration solutions were prepared by diluting his mix in the pooled sample with methanol and water added to match the composition of the samples. The concentrations prepared included 0.25, 0.5, 2.5, 5, 12.5, 25, 50, 125, 250, and 500 ng/mL. Separate hydrophilic interaction liquid chromatography (HILIC) methods were developed to separate metabolites in negative and positive ionization modes. For negative mode, a 2.1 × 100 mm, 2.7 μm Infinity Lab Poroshell HILIC-Z, P column (Agilent Technologies) was used. A mobile phase at pH 9 was prepared for gradient elution with 10 mM ammonium acetate in water (A) and 10 mM ammonium acetate in 90% acetonitrile (B) and pH adjusted with ammonium hydroxide. The additive medronic acid (5 mM methylenediphosphonic acid) was added to A and B at final 5 μM concentration to improve peak shape (72). A sample volume of 5 μL was injected onto the column held at 30 °C, and eluted at a flow rate of 0.3 mL/min with the following gradient: hold at 90% B from 0 to 1.7 min, 90 to 60% from 1.7 to 10 min, hold at 60% B from 10.2 to 12.7 min, return to 90% B at 13.5 min, and equilibrate with 90% B till 20 min. For positive ion mode analysis, a 2.1 × 100 mm, 1.7 um Acquity BEH Amide column (Waters Corporation) was used. A mobile phase at pH 10 was prepared and composed of 0.1% ammonium hydroxide in Milli-Q water (A) and 0.1% ammonium hydroxide in 90% acetonitrile. A 5 μL sample volume was injected onto the column held at 40 °C, and eluted at 0.3 mL/min with the same gradient as above. The autosampler was washed between injections to prevent carryover with two successive wash solutions, 50:50 acetonitrile/water followed by 95:5 acetonitrile/water.

To quantify metabolites, we used liquid chromatography (Accela Open Autosampler and Accela 1250 Pump, Thermo Scientific) coupled to a heated electrospray ionization source (H- ESI) and a triple quadrupole mass spectrometer (TSQ Vantage, Thermo Scientific) operated under selected reaction monitoring (SRM) mode. Optimal SRM parameters (s-lens, collision energy, fragment ions) for each target compound were determined with authentic standards purchased from Sigma. The complete list of metabolites is provided in Table S5. We monitored two SRM transitions per compound for quantification and confirmation and generated external calibration curves based on peak area for each compound. We converted raw data files from proprietary Thermo (.RAW) format to mzML using the msConvert tool(73) prior to processing with El-MAVEN(74). Samples were analyzed in random order within each batch, and the pooled QC sample was analyzed every six injections.

### Phylogenetic analysis

Sequences were collected from the Genbank database(75) for the following chitin degradation pathway proteins: The N-terminal region of ChiA/ChiA-like, the C-terminal region of ChiA/ChiA-like, UgpA, UgpE, NagZ, NagK, NagA, and NagB. Orthologs found within *Prochlorococcus* MIT1303 were used as protein search queries using BLASTP(76), with the top 500 or 250 hits recovered in each case. Each set of sequences were then aligned in MAFFT(77) with the automatic algorithm selection option. Aligned sequences were then used for phylogenetic reconstruction using IQTree(78) with automatic best-fitting model selection. All sequence alignment and phylogenetic data files are available in SI data (Data S1, SI Text 1 files), with alignment and tree filenames in each case describing the algorithms and parameters used for these reconstructions. Several BLAST hits for the ChiA and ChiA- like genes in SynPro overlapped, with some protein sequences containing multiple domains homologous to different chitinase orthologs in other bacteria. A phylogenomic analysis of our alignment data showed that the SynPro variant was likely the result of a fusion between genes from Planctomycetes, before or after horizontal gene transfer into SynPro (see SI Text 1 and SI Text 1 files for a detailed analysis of the protein fusion history).

## Acknowledgments

We thank members of the Chisholm lab for thoughtful discussions, and Nikolai Radzinski for insightful discussions and useful references regarding microbial aggregation. This reseach was supported by grants from the Simons Foundation (Award ID #509034SCFY20 to R.B. and S.W.C., and SCOPE Award ID 329108 to M.J. Follows; Life Sciences Award IDs 337262, 647135, 736564 to S.W.C.; SCOPE Award ID 329108 and 721246 to S.W.C.). G.C. was also supported by the EMBO Long-Term Fellowship (ALTF 904-2018) and by the Human Frontier Science Program (LT000069/2019-L). This paper is a contribution from the Simons Collaboration on Ocean Processes and Ecology (SCOPE).

## Contributions

G.C., R.B., A.Y. and S.W.C. designed experiments.

G.C., J.S., X.L. and A.M. performed experiments.

T.H., E.T. and R.B. performed bioinformatic analyses.

K.L., M.K.S., G.S. and E.B.K. performed metabolite analyses.

G.F. and J.P. performed phylogenetic analyses.

G.C, R.B. and S.W.C. interpreted data.

R.B. developed macroevolutionary model.

R.B., G.C., and S.W.C. wrote the paper with contributions from all authors.

## Notes

### Competing Interest Statement

The authors have declared no competing interest.

### Summary of Updates

Added contact information of the corresponding authors Giovanna Capovilla Rogier Braakman Sallie W. Chisholm

