## Supplementary material for "Chitin utilization by marine picocyanobacteria and the evolution of a planktonic lifestyle": Capovilla_Braakman_Supplemental.pdf

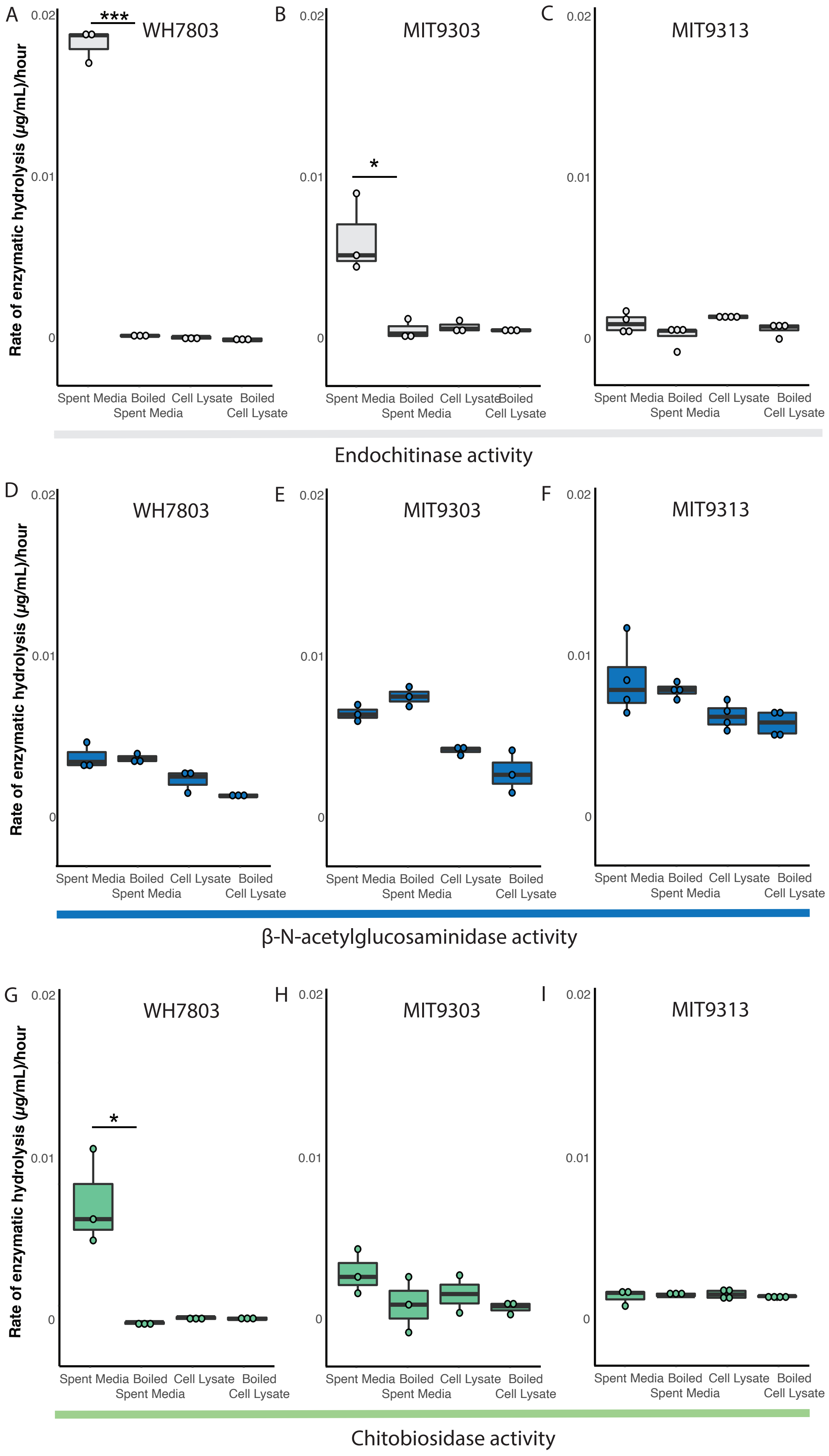

A
WH7803
***
B
MIT9303
*
C
MIT9313
Rate of enzymatic hydrolysis (µg/mL)/hour
0.02
0.01
0
Spent Media
Boiled Spent Media
Cell Lysate
Boiled Cell Lysate
Endochitinase activity
D
WH7803
E
MIT9303
F
MIT9313
β-N-acetylglucosaminidase activity
G
WH7803
*
H
MIT9303
I
MIT9313
Chitobiosidase activity

**FIGURE S1. Endochitinase and Exochitinase activities in marine picocyanobacteria**. Chitinase activity in *Synechococcus* WH7803, *Prochlorococcus* MIT9303 and MIT9313 were measured in the spent media and cell lysates of cultures growing in Pro99 (Sea water media) amended with chitosan to a final concentration of 56 µg/ml. The degradation of chitin oligomers attached to the fluorophore methylumbelliferyl is estimated by the luminescence intensity. Three activities are shown. A-C) The endochitinase activity (ability to cleave glycosidic bonds at internal sites of the chitin polymer) shown in grey, D-F) β-N-acetylglucosaminidase activity (ability to cleave GlcNAc monomers from the nonreducing terminal of the chitin polymer) shown in blue G-I) Chitobiosidase activity (ability to cleave dimeric units of GlcNAc from the nonreducing terminal of the chitin polymer) shown in green. Statistical significance has been performed between each sample and its control with a Welch t-test. Data in (a), (b) and (c) are also shown in Figure 2.

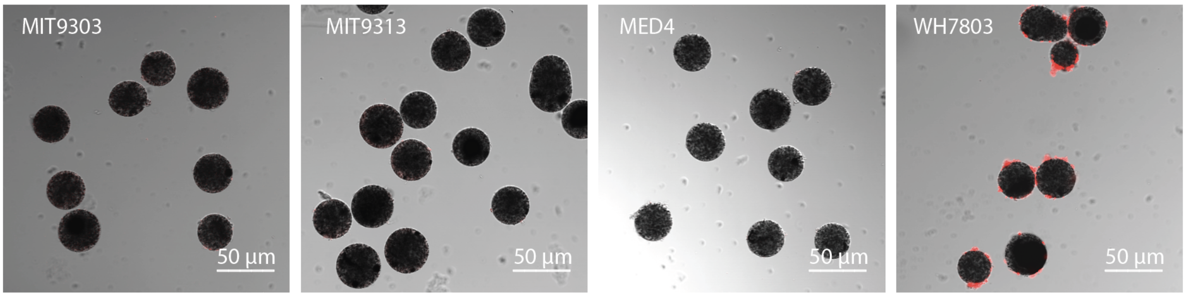


**FIGURE S2. Colonization of agarose particles.** Images of magnetic agarose beads analyzed in Figure 3 to illustrate consistency and uniformity across beads. Beads are shown in bright field mode. *Synechococcus* and *Prochlorococcus* are detected by their autofluorescence and highlighted in red. White bars are 50 µm long

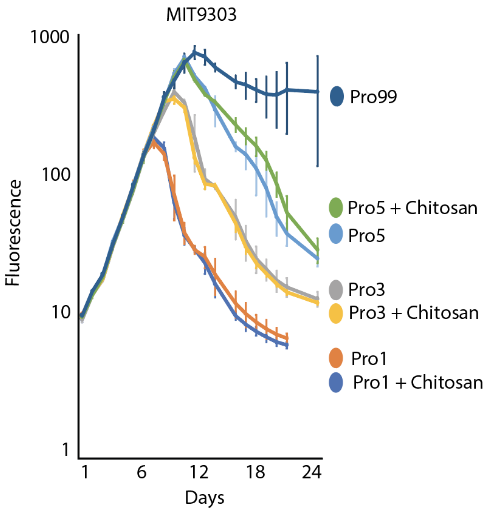


**FIGURE S3. Response of *Prochlorococcus* to chitosan under nitrogen stress.** Growth of *Prochlorococcus* MIT9303 in continuous light (at 12 µmol photons $m^{-2}$ $s^{-1}$) in a variety of different media, monitored by relative bulk culture chlorophyll fluorescence. A. Different colors show the growth patterns of triplicate cultures grown in Pro99 media with different concentrations of $NH_4^+$, the sole N-source, with and without added chitosan (56 µg/ml). The concentrations of $NH_4^+$ were 800 µM in Pro99, 250 µM in Pro5, 150 µM in Pro3, and 50 µM in Pro1.

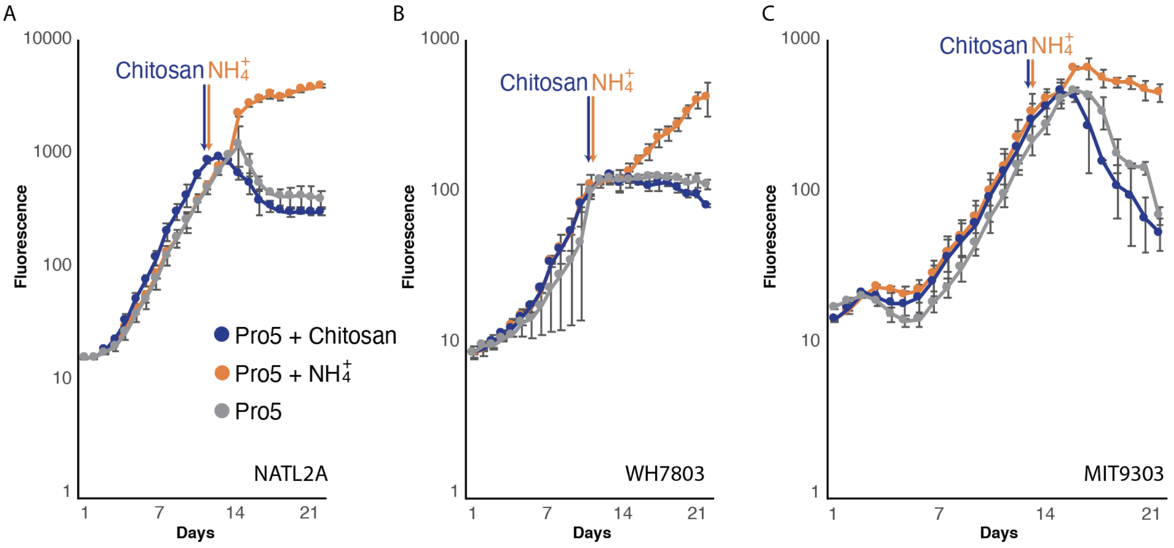


**FIGURE S4. Comparison of the response to chitosan and ammonia under nitrogen stress.** Growth of *Prochlorococcus* NATL2A, *Synechococcus* WH7803, and *Prochlorococcus* MIT9303 in N-depleted media (denoted as "Pro5" to indicate the lowered nitrogen-to-phosphorus ratio of N:P = 5) amended with $NH_4^+$ or chitosan when cells reached N-starved stationary phase. Different colors show the growth patterns in media Pro5 that has been amended with $NH_4^+$ (orange) at the time of the arrow, to a final concentration of 800 µM (N:P =16), or chitosan (blue) to a final concentration of 56 µg/ml, or non-amended (grey). Cultures were grown in continuous light (at 15 µmol photons $m^{-2}$ $s^{-1}$) and growth was monitored by relative bulk culture chlorophyll fluorescence.

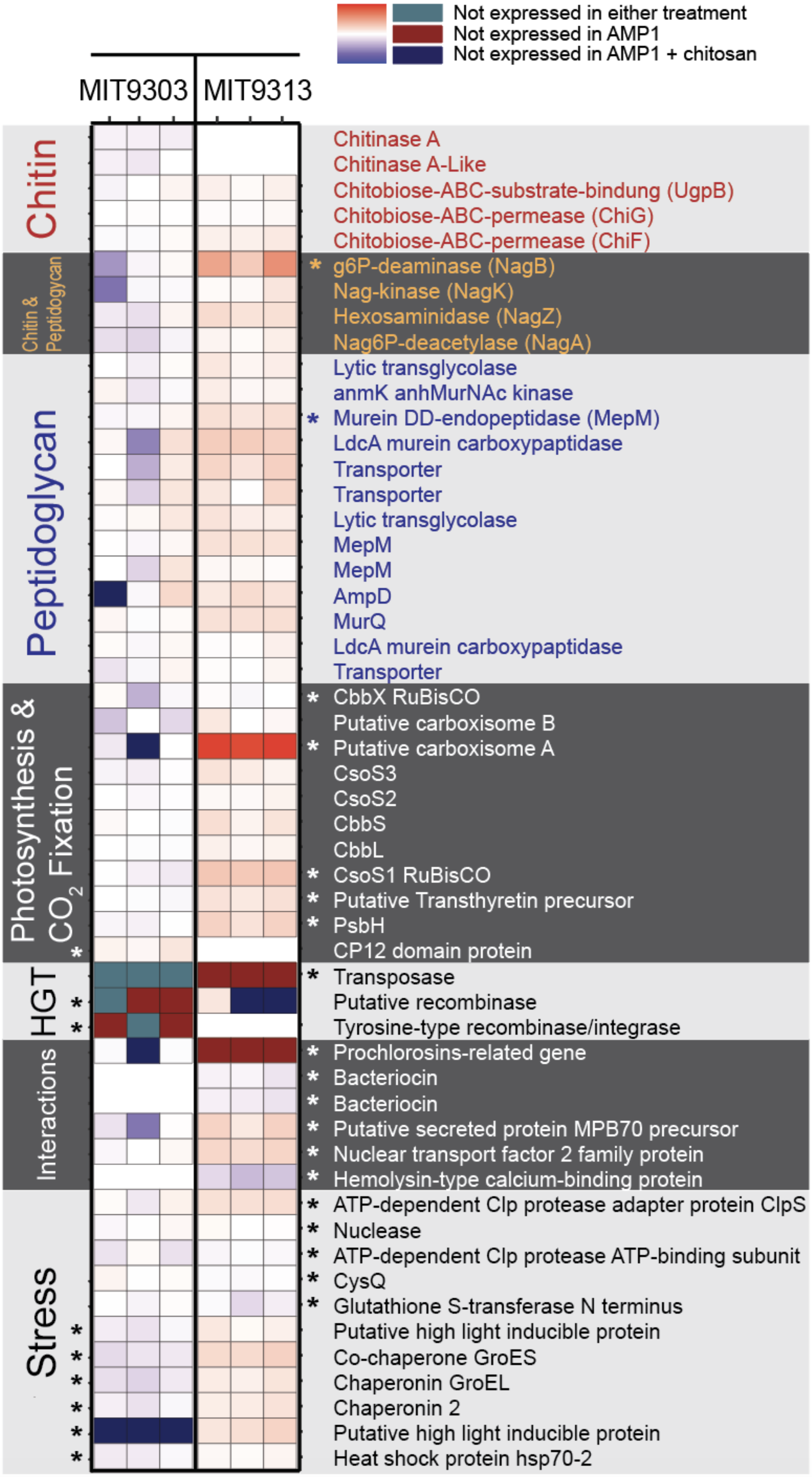
Not expressed in either treatment
Not expressed in AMP1
Not expressed in AMP1 + chitosan
MIT9303
MIT9313
Chitin
Chitin & Peptidoglycan
Peptidoglycan
Photosynthesis & CO2 Fixation
HGT
Interactions
Stress
Chitinase A
Chitinase A-Like
Chitobiose-ABC-substrate-bindung (UgpB)
Chitobiose-ABC-permease (ChiG)
Chitobiose-ABC-permease (ChiF)
* g6P-deaminase (NagB)
Nag-kinase (NagK)
Hexosaminidase (NagZ)
Nag6P-deacetylase (NagA)
Lytic transglycolase
anmK anhMurNAc kinase
* Murein DD-endopeptidase (MepM)
LdcA murein carboxypaptidase
Transporter
Transporter
Lytic transglycolase
MepM
MepM
AmpD
MurQ
LdcA murein carboxypaptidase
Transporter
* CbbX RuBisCO
Putative carboxisome B
* Putative carboxisome A
CsoS3
CsoS2
CbbS
CbbL
* CsoS1 RuBisCO
* Putative Transthyretin precursor
* PsbH
CP12 domain protein
* Transposase
Putative recombinase
Tyrosine-type recombinase/integrase
* Prochlorosins-related gene
* Bacteriocin
* Bacteriocin
* Putative secreted protein MPB70 precursor
* Nuclear transport factor 2 family protein
* Hemolysin-type calcium-binding protein
* ATP-dependent Clp protease adapter protein ClpS
* Nuclease
* ATP-dependent Clp protease ATP-binding subunit
* CysQ
* Glutathione S-transferase N terminus
Putative high light inducible protein
Co-chaperone GroES
Chaperonin GroEL
Chaperonin 2
Putative high light inducible protein
Heat shock protein hsp70-2

**FIGURE S5. Quantitative analysis of gene expression in *Prochlorococcus* in response to chitosan addition.** Heat map representing the relative expression of genes in the two *Prochlorococcus* strains 24 hours after addition to chitosan, reported as a qualitative representation in figure 5B. Three biological replicates are shown for each strain. Transcriptional enrichment between the samples amended with chitosan and those that were not is indicated in shades of red, while transcriptional depletion is represented in shades of blue. Those genes which were statistically significantly differentially expressed are marked with a black or white star. All genes in the chitin degradation and peptidoglycan recycling pathways, as well as all genes of the $CO_2$-fixation, are shown. Genes are grouped by functional categories and represented in shades of grey.

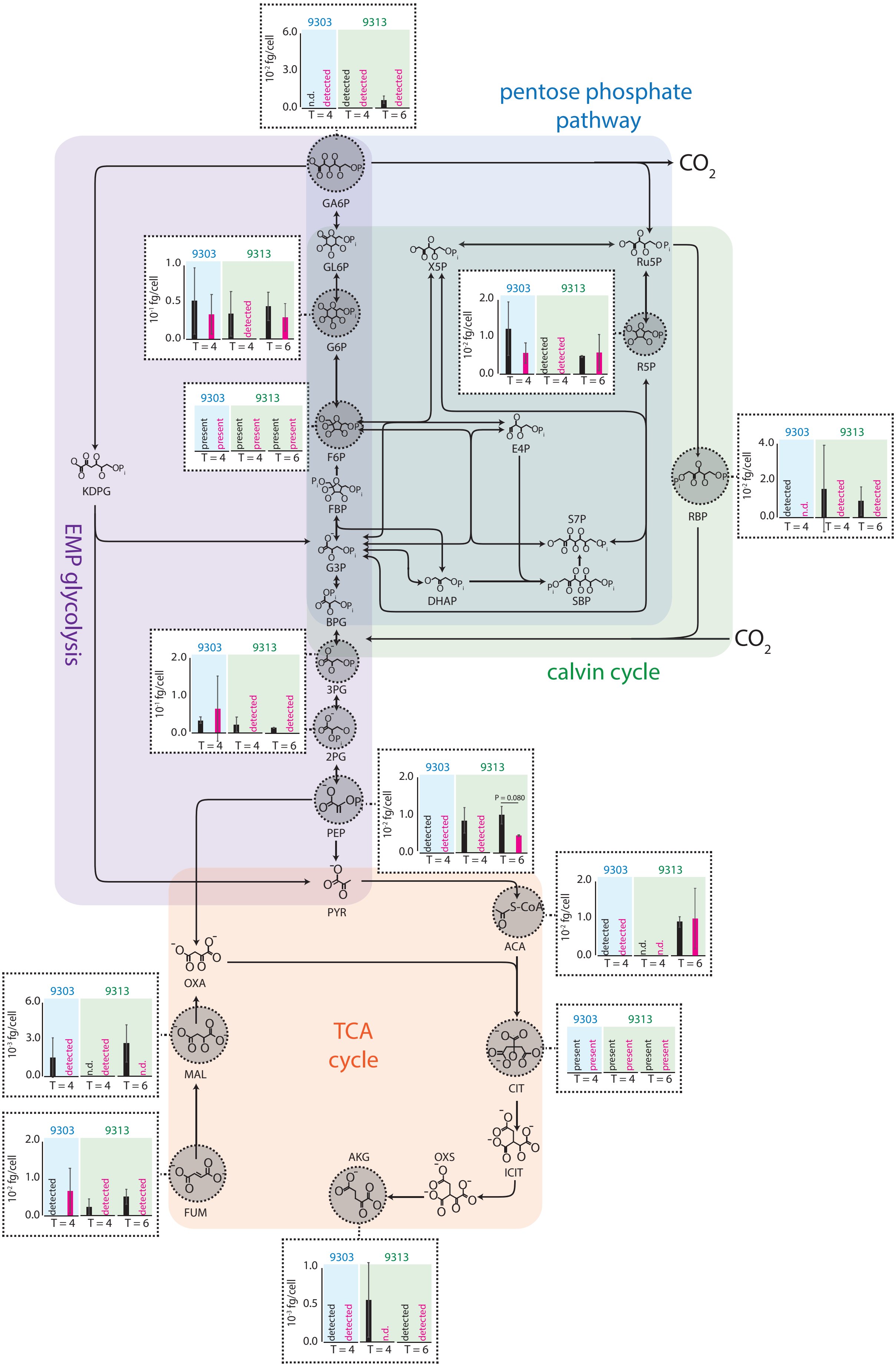

pentose phosphate pathway
EMP glycolysis
calvin cycle
TCA cycle
CO2
GA6P
GL6P
G6P
F6P
FBP
G3P
BPG
3PG
2PG
PEP
PYR
KDPG
X5P
Ru5P
R5P
E4P
S7P
SBP
DHAP
RBP
ACA
OXA
MAL
FUM
CIT
ICIT
OXS
AKG
9303
9313
10⁻² fg/cell
10⁻¹ fg/cell
10⁻³ fg/cell
n.d.
detected
present
T = 4
T = 6
P = 0.080

**FIGURE S6. Metabolomic analysis of core carbohydrate pathways in response to chitosan addition in *Prochlorococcus*.** Concentrations, in femtograms per cell, of intermediates of core carbohydrate metabolic pathways occurring downstream of the chitin degradation pathway shown in Figure 5C, on days 4 and 6 after chitosan additions (pink) compared to unamended controls (black). Major pathways within the metabolic core are highlighted in different colors. Error bars show the standard deviation of 3 biological replicates of MIT9303 and MIT9313 on day 4, and of 2 replicates of MIT9313 on day 6. Samples in which metabolites were observed in only a single replicate are denoted ‘detected’, while samples, where metabolites were not observed in any replicates, are labeled ‘n.d.’ (not detected). Fructose-6P (F6P) and citrate (CIT) were present in most samples, but their levels could not be quantified due to interference by the organic carbon matrix. Abbreviations: GA6P - Gluconate-6-phosphate, GL6P - Gluconolactone-6-phosphate, G6P - Glucose-6-phosphate, F6P - Fructose-6-phosphate, FBP - Fructose bisphosphate, G3P - Glyceraldehyde-3-phosphate, DHAP - Dihydroxyacetone-phosphate, SBP - Sedoheptulose bisphosphate, S7P - Sedoheptulose-7-phosphate, E4P - Erythrose-4-phosphate, X5P - Xylulose-5-phosphate, R5P - Ribose-5-phosphate, Ru5P - Ribulose-5-phosphate, RBP - Ribulose bisphosphate, KDGP - 2-Keto-3-deoxy-6-phosphogluconate, BPG - bisphospho-glycerate, 3PG - 3-phosphoglycerate, 2PG - 2-phosphoglycerate, PEP - Phosphoenoylpyruvate, PYR - Pyruvate, ACA - Acetyl-CoA, OXA - Oxaloacetate, MAL - Malate, FUM - Fumarate, CIT - Citrate, ICIT - Isocitrate, OXS - Oxalosuccinate, AKG - alpha-ketoglutarate.

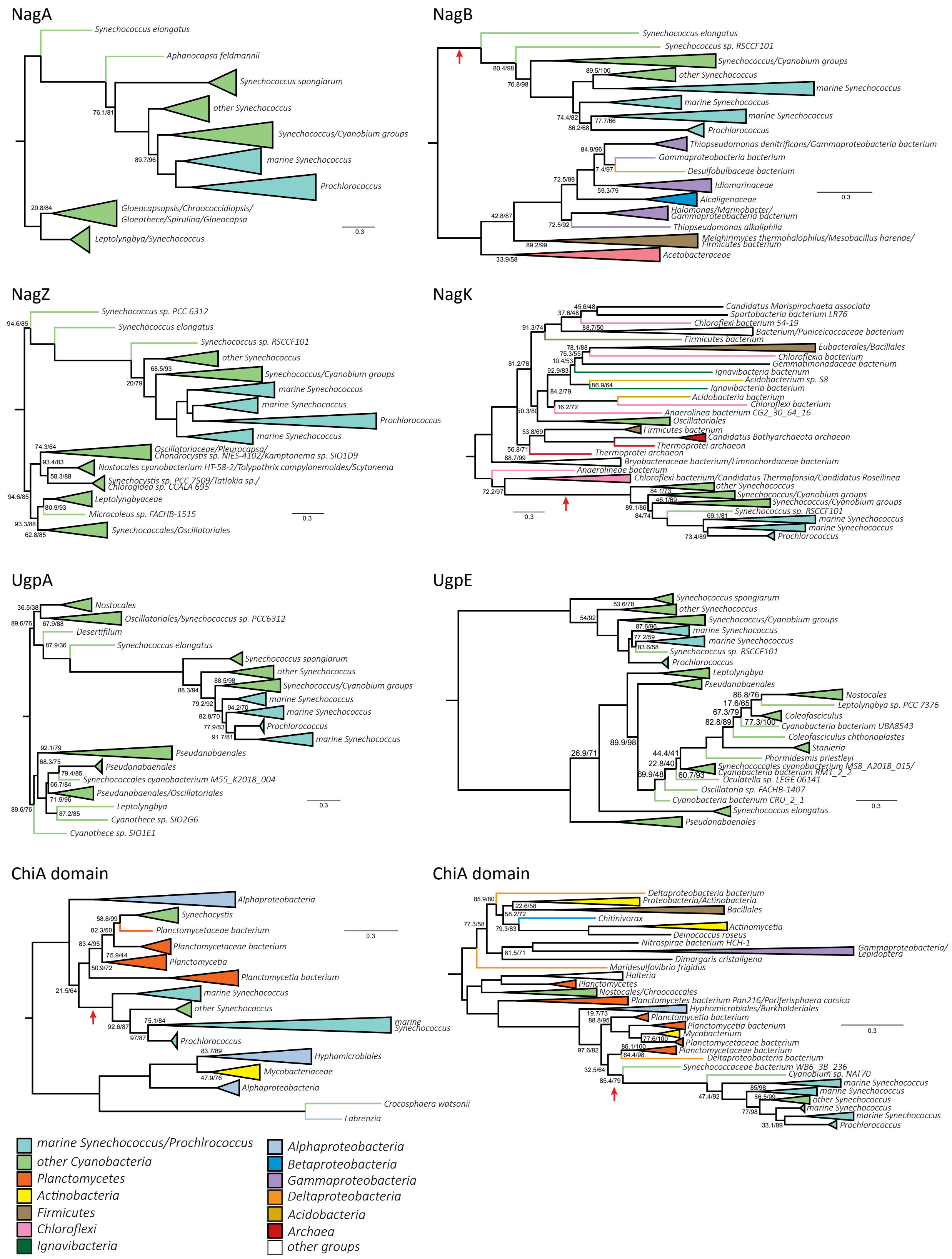


**FIGURE S7. Phylogenies of genes involved in chitin utilization in marine picocyanobacteria.** Red arrows indicate inferred HGTs into SynPro ancestor lineages. Branch lengths are indicated by included scale bars (average substitutions/site). High-level taxonomic identities are indicated by color coding, with specific represented groups labeled for terminal and collapsed groups. In each case, trees are rooted at the branch closest to the midpoint that preserves the monophyly of Synechococcales groups including SynPro. Branch supports (approximate likelihood ratio test/bootstrap value) are included for bipartitions with less than 90% support for either metric. Phylogenies of the two ChiA domains and NagK are also shown in figure 6.

**SI Text 1 and SI Text 1 files**: Phylogenomic analysis of Chitinase Domains within SynPro proteins ChiA and ChiA-like.

**Data S1**: Chitinase pathway final alignment and tree files.

**Table S1**. Annotated genes

**Table S2**. List of primers used for the qPCR experiment

| **Gene** | **Forward primer** | **Reverse primer** |
|---|---|---|
| chitinase A (ChiA) | GGTGCAAGACATCAATGTCG | GCAGCCCAAGAATCAAAAAG |
| chitinaseA-like (ChiA-like) | GTTCTCGTTGAGCAAAGCCAG | ACCCCACCAAAGATCACCAG |
| chitobiose-ABC-permease (UgpA) | CCAAAACAGGTCTCGACGTT | AACATCGGATCACCAGCAAG |
| chitobiose-ABC-permease (UgpE) | CCATTGCTTTGGCTGGTAAG | TCGGTTCGGCAGGTAGTAAG |
| chitobiose-ABC-substrate-binding (UgpB) | ACGTTGGCAGGATGTACCTT | GAATCGTCAGGGACCACAGT |
| g6P-deaminase (NagB) | GGCAGCTGATCGGCGCAGC | GAAGATGCAACTGCTGCGGG |
| hexosaminidase (NagZ) | CGACCCAGATCAGCCTTTAG | TTCCCGCGTAAGAAAAGTTG |
| nag6P-deacetylase (NagA) | GAGCGGGTGCTCAGTGTTC | AGCCCTCCATTGATTTGTAGA |
| nag-kinase (NagK) | GGAGCAAGCGGAATAGAACA | ATCCCCAGTTGCTAGGCATT |
| rnpB | TGAGGAGAGTGCCACAGAAA | ACCTCTCGATGCTGCTGGT |

**Table S3**
RNA-Seq data

**Table S4**
Metabolites data

**Table S5**
Complete set of metabolites measured using liquid chromatography-mass spectrometry (LC-MS). A novel method was developed to target this set of metabolites. The polarity (+ or -, for positive or negative) indicates the ionization mode used to analyze the metabolite. For select metabolites, concentrations could not be determined due to interference by the organic carbon matrix, these are marked as 'presence / absence' in the table.

| **compound** | **KEGG** | **polarity** | **Detection in *Prochlorococcus*** |
|---|---|---|---|
| 2-keto-3-deoxy-6-phosphogluconic acid | C04442 | - | not detected |
| 6-phosphogluconic acid | C00345 | - | yes |
| acetyl coenzyme A | C00024 | - | yes |
| alpha-ketoglutaric acid | C00026 | - | yes |
| chitobiose | C01674 | + | yes |
| citric acid | C00158 | - | presence / absence |
| 3- and 2-phosphoglyceric acid | C00197 / C00631 | - | yes, but cannot be analytically separated |
| D-glucosamine | C00329 | + | yes |
| D-glucosamine 6-phosphate | C00352 | - | yes |
| D-Ribose 5-phosphate | C00117 | - | yes |
| D-ribulose 1,5-bisphosphate | C01182 | - | yes |

| | | | |
|---|---|---|---|
| fructose 6-phosphate | C00085 | - | presence / absence |
| fumaric acid | C00122 | - | yes |
| glucose 6-phosphate | C00092 | - | yes |
| malic acid | C00149 | - | yes |
| N-Acetyl-D-glucosamine 6-phosphate | C00357 | - | yes |
| phosphoenolpyruvate | C00074 | - | yes |
